# Investigation of the Stress and Sleep Physiology Correlates of Next-Day Memory for Details of a Social Stressor Testing Environment

**DOI:** 10.1101/2021.01.28.428506

**Authors:** Ryan Bottary, Sarah M. Kark, Ryan T. Daley, Dan Denis, Tony J. Cunningham, Jessica D. Payne, Elizabeth A. Kensinger

## Abstract

Despite evidence which demonstrates that psychosocial stress interacts with sleep to modulate memory, research that has examined next-day memory for the stressful environment itself has not accounted for post-stressor sleep. Here, participants completed the Trier Social Stress Test or a matched control task with psychophysiological monitoring and stress hormone assays. After a 24-hour delay that included overnight polysomnographically-recorded sleep, memory for objects in the testing room was assessed by having participants draw the testing room from the previous day from memory. As expected, stressed participants mounted greater psychophysiological and stress hormone responses to the stressor than participants in the control condition. However, there was only weak evidence that stress reactivity and post-encoding sleep interacted to modulate memory for testing room details. Instead, NREM sleep physiology on the night following testing room encoding was positively associated with memory for testing room details, though this association occurred in the control, but not stressed, participants.

## Introduction

Human episodic memories are influenced by acute stress. However, memory effects vary as a function of stressor timing relative to memory encoding and the degree to which learned material is related to the stressor itself (for review see Shields et al., 2017). Findings from a recent meta-analysis suggest that acute stress improves long-term retention of encoded material if (1) the stressor occurs in close temporal proximity to encoding and (2) the learned material is directly related to the stressor (Shields et al., 2017). Interestingly, a series of studies (Herten, Otto, et al., 2017; Herten, Pomrehn, et al., 2017; Wiemers et al., 2013, 2014) that probed memory during a psychosocial evaluative stressor that included performance judges (i.e. the Trier Social Stress Test; Kirschbaum et al., 1993) found that stressed participants preferentially remembered objects central to, compared to those peripheral to, this stressful experience. While these studies systematically investigated several potential task paradigm-dependent memory modulators (e.g. impacts of attitudes and affect of the judges, measures of attention, encoding-retrieval delay interval), it remains unclear if individual differences in task-dependent stress reactivity relate to observed memory patterns. Further, three out of these four above mentioned studies (Herten, Otto, et al., 2017; Wiemers et al., 2013, 2014) tested memory at least 24 hours after memory encoding, an interval likely to include variable amounts of overnight sleep, but they did not specifically measure sleep. Prior work suggests that individual differences in stress responsivity at encoding, measured by cortisol, skin conductance and heart rate, interacts with post-encoding sleep to selectively strengthen memory (Bennion et al., 2015; Cunningham et al., 2014; Kim et al., 2019). Further, electroencephalographic rhythms during rapid eye-movement (REM; Hutchison & Rathore, 2015; Kim et al., 2019) and non-REM sleep (Latchoumane et al., 2017; Lehmann et al., 2016) predict retention of emotional memories. Post-encoding REM sleep theta power, in particular, has recently been shown to correlate with the retrieval of emotional information encountered *after* the same TSST stressor used here (Kim et al., 2019). The present study examines how retrieval of information encountered *during* that TSST stressor is influenced by variation in stress responding to the TSST as well as variability in post-stressor sleep physiology.

We aimed to address this question by exploring the interactive effects of stress responsivity and post-encoding sleep physiology on memory for objects in the testing environment during the TSST (Stress Condition) or a matched control task (Control Condition). For participants in the Stress Condition, central objects were classified as those proximal to the judges (located on a table at which judges sat during evaluation) and objects critical for evaluation (e.g. microphone, camcorder, stopwatch, psychophysiology recording equipment). We hypothesized that individual differences in stress responsivity, measured by stress hormone assays and psychophysiology, would positively interact with post-encoding sleep physiology to selectively enhance memory for central objects. To assess general effects of sleep physiology on memory without a salient stressor, we conducted these analyses for participants in the Control Condition, who completed the same tasks as those in the Stress Condition, though without judges and evaluative objects (e.g. microphone, camcorder, stopwatch) present. Evaluative objects were removed from the testing environment in the Control Condition to reduce potential stress induction confounds.

## Materials and Methods

### Participants

Sixty-five healthy, right-handed, native English-speaking young adult participants (age 18-31 years; 35 female;) were recruited from Boston College and the greater Boston area as part of a larger study examining the effects of sleep and stress on emotional memory (see **Table 1** for participant demographics). Participants had normal or corrected-to-normal vision, reported no history of chronic medical conditions, neurological, psychiatric or sleep disorders, or current use of psychoactive medications. Seven participants (4 female, 3 male) did not complete the memory test and were therefore excluded from analyses. Study procedures were approved by the Boston College Institutional Review Board. Participants provided informed consent prior to study participation and were compensated for their time.

**Table 1.**
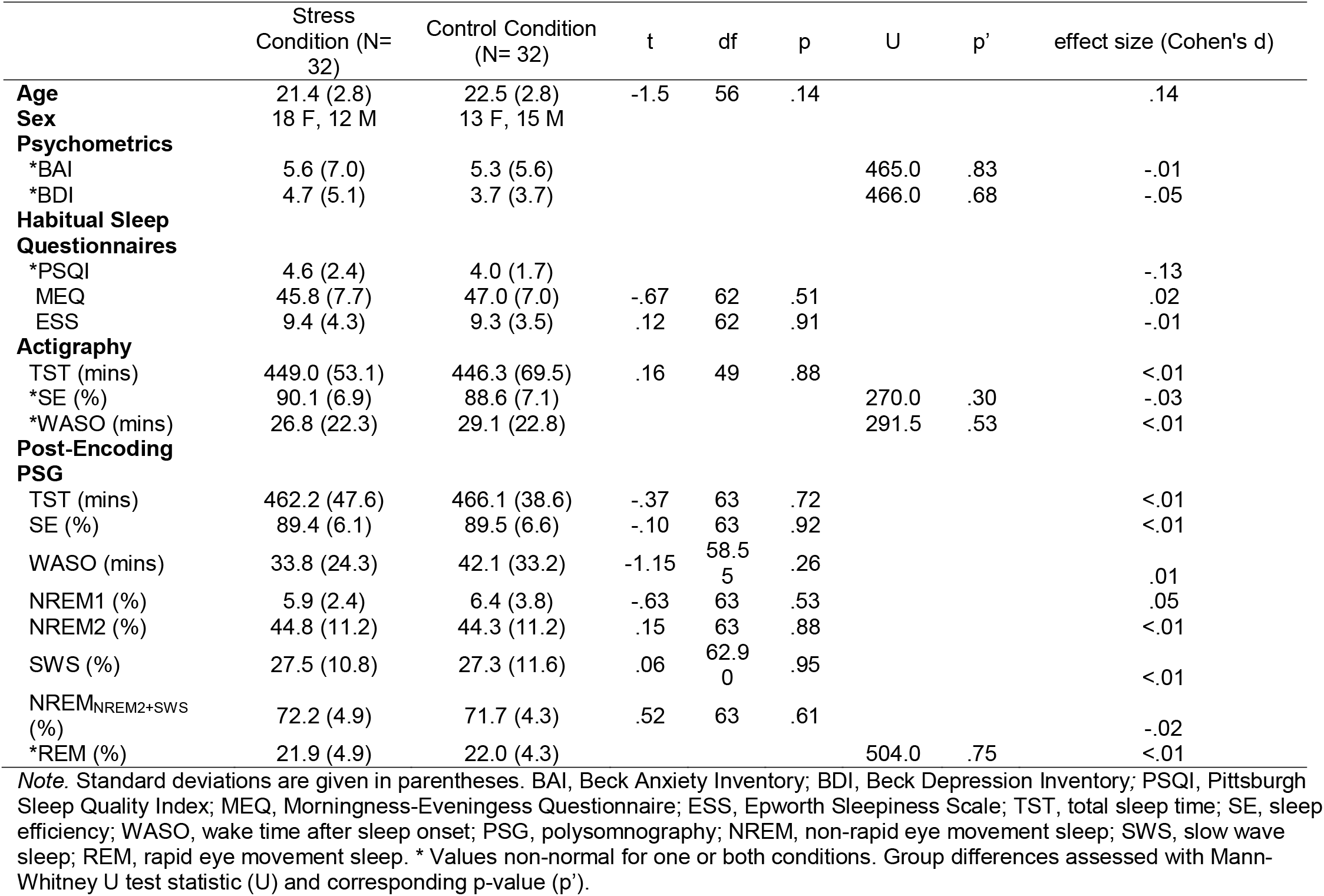
Participant demographic and psychometric and sleep variable values.

|  | Stress<br>Condition (N=32) | Control Condition<br>(N= 32) | t | df | p | U | p' | effect size (Cohen's d) |
| --- | --- | --- | --- | --- | --- | --- | --- | --- |
| **Age** | 21.4 (2.8) | 22.5 (2.8) | -1.5 | 56 | .14 |  |  | .14 |
| **Sex** | 18 F, 12 M | 13 F, 15 M |  |  |  |  |  |  |
| **Psychometrics** |  |  |  |  |  |  |  |  |
| *BAI | 5.6 (7.0) | 5.3 (5.6) |  |  |  | 465.0 | .83 | -.01 |
| *BDI | 4.7 (5.1) | 3.7 (3.7) |  |  |  | 466.0 | .68 | -.05 |
| **Habitual Sleep Questionnaires** |  |  |  |  |  |  |  |  |
| *PSQI | 4.6 (2.4) | 4.0 (1.7) |  |  |  |  |  | -.13 |
| MEQ | 45.8 (7.7) | 47.0 (7.0) | -.67 | 62 | .51 |  |  | .02 |
| ESS | 9.4 (4.3) | 9.3 (3.5) | .12 | 62 | .91 |  |  | -.01 |
| **Actigraphy** |  |  |  |  |  |  |  |  |
| TST (mins) | 449.0 (53.1) | 446.3 (69.5) | .16 | 49 | .88 |  |  | <.01 |
| *SE (%) | 90.1 (6.9) | 88.6 (7.1) |  |  |  | 270.0 | .30 | -.03 |
| *WASO (mins) | 26.8 (22.3) | 29.1 (22.8) |  |  |  | 291.5 | .53 | <.01 |
| **Post-Encoding PSG** |  |  |  |  |  |  |  |  |
| TST (mins) | 462.2 (47.6) | 466.1 (38.6) | -.37 | 63 | .72 |  |  | <.01 |
| SE (%) | 89.4 (6.1) | 89.5 (6.6) | -.10 | 63 | .92 |  |  | <.01 |
| WASO (mins) | 33.8 (24.3) | 42.1 (33.2) | -1.15 | 58.5<br>5 | .26 |  |  | .01 |
| NREM1 (%) | 5.9 (2.4) | 6.4 (3.8) | -.63 | 63 | .53 |  |  | .05 |
| NREM2 (%) | 44.8 (11.2) | 44.3 (11.2) | .15 | 63 | .88 |  |  | <.01 |
| SWS (%) | 27.5 (10.8) | 27.3 (11.6) | .06 | 62.9<br>0 | .95 |  |  | <.01 |
| NREM<sub>NREM2+SWS</sub> (%) | 72.2 (4.9) | 71.7 (4.3) | .52 | 63 | .61 |  |  | -.02 |
| *REM (%) | 21.9 (4.9) | 22.0 (4.3) |  |  |  | 504.0 | .75 | <.01 |

*Note.* Standard deviations are given in parentheses. BAI, Beck Anxiety Inventory; BDI, Beck Depression Inventory; PSQI, Pittsburgh Sleep Quality Index; MEQ, Morningness-Eveningness Questionnaire; ESS, Epworth Sleepiness Scale; TST, total sleep time; SE, sleep efficiency; WASO, wake time after sleep onset; PSG, polysomnography; NREM, non-rapid eye movement sleep; SWS, slow wave sleep; REM, rapid eye movement sleep. \* Values non-normal for one or both conditions. Group differences assessed with Mann-Whitney U test statistic (U) and corresponding p-value (p').

### Pre-Study Sleep Monitoring

In the three days prior to testing, participants were asked to keep a regular sleep schedule, aiming for a bedtime prior to 2 AM and sleeping for at least 7 hrs. To enhance compliance, participants were monitored with wrist actigraphy and daily online sleep logs. Participants were required to abstain from caffeine, alcohol, and tobacco for the 24 hours prior to the start of the study and for the duration of the study. Prior to testing, participants completed habitual sleep questionnaires including the Pittsburgh Sleep Quality Index (PSQI; Buysse et al., 1989), Morningness-Eveningness Questionnaire (MEQ; Horne & Östberg, 1976) and Epworth Sleepiness Scale (ESS; Johns, 1991) as well as questionnaires about depression (Beck Depression Inventory II; BDI-II; Beck et al., 1996) and anxiety (Beck Anxiety Inventory; BAI; Steer & Beck, 1997) (for descriptive statistics see **Table 1**).

### Stress Manipulation

Participants were randomly assigned to either the Stress Condition, in which they were subject to a psychosocial stress protocol (i.e. the TSST), or a matched control task. All participants were provided note-taking materials, then given 10 mins to prepare a 5-min speech on a topic (e.g., explain why they are the best candidate for a job position) using only truthful information about themselves. Participants in the Stress Condition were informed that their speech would be evaluated for verbal and nonverbal performance by two judges and that their performance would be video- and audio-recorded for later analysis. Participants in the Control Condition were informed of their condition designation to reduce anticipatory anxiety. After 10 mins had elapsed, participants were escorted to a separate room with two seated judges (or an empty room in the Control Condition). Judges maintained a neutral expression throughout testing. All participants were outfitted with psychophysiological measurement electrodes that continuously recorded skin conductance level and heart rate. In the Stress Condition, participants’ notes were abruptly taken from them prior to the speech and they were asked to give their speech from memory. If their speech ended in under 5 mins, judges instructed participants to continue talking until the full 5-mins were up. Immediately following the speech, participants were instructed to perform an arithmetic task aloud for 5 min (e.g., “Continuously subtract 13 starting from the number 1022 as quickly and accurately as possible”). If Stress participants made a mistake, judges instructed them to start over. Participants in the Control Condition were instructed to read their speech aloud from their notes and completed a simpler version of the arithmetic task while alone in the room.

### Incidental Encoding

All objects located in the testing environment served as targets for incidental encoding (see **Figure 1** for photographs of the testing environment). The testing environment was set-up in the same way for all participants with the exception that the microphone and camcorder were not present for Control participants. This decision was made to avoid the potential confound of inducing evaluation stress in the Control participants. Additionally, judge’s materials (i.e. clipboards, pens and stopwatch) were not present for Control participants.

**Figure 1.**
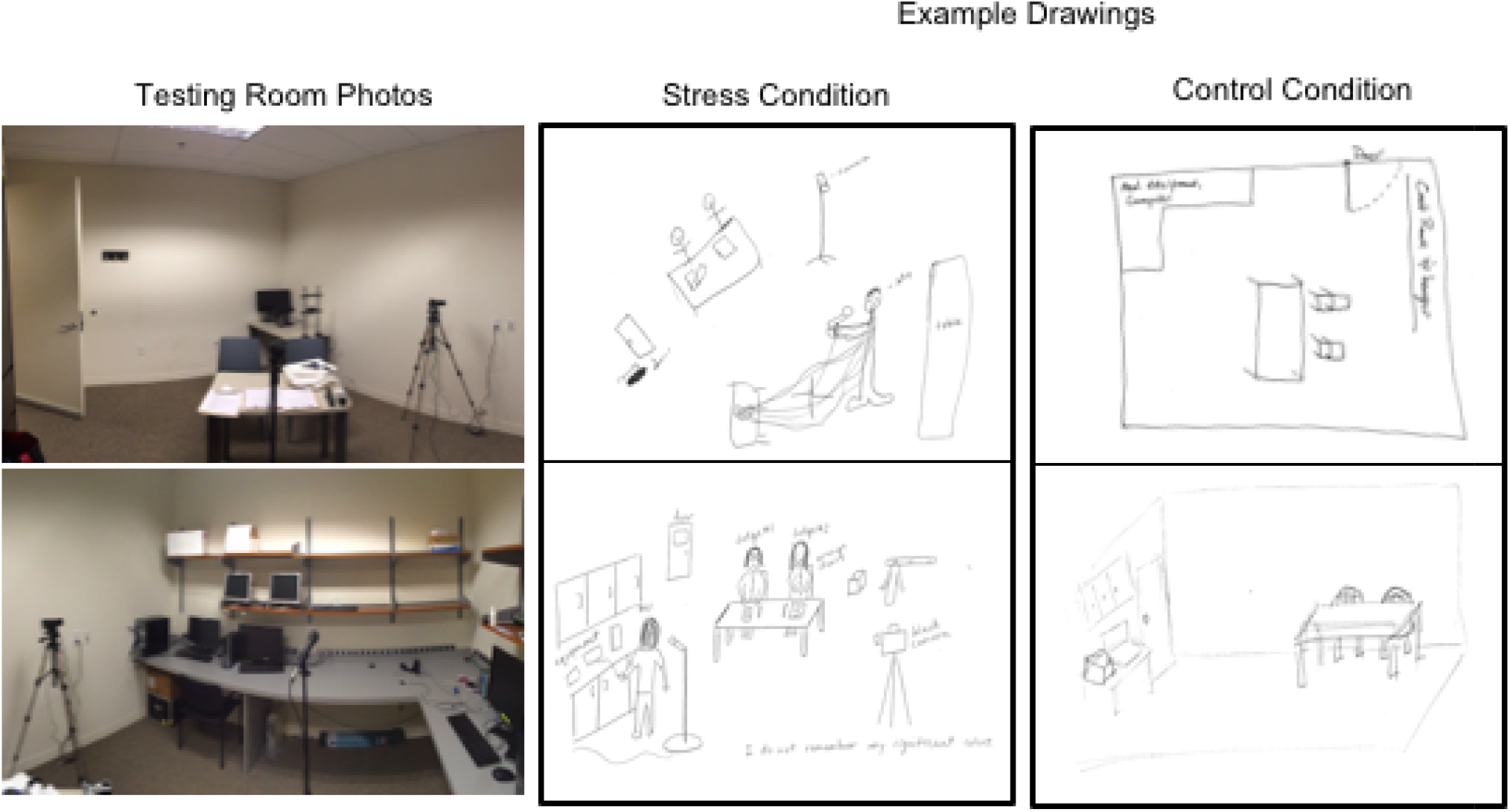
Photographs of the testing room and example drawings. The photographs on the far left depict the testing room from (top) the perspective of the participant during testing as well as (bottom) the participant’s surroundings. The center block contains two example drawings from participants that underwent the TSST. The top drawing received a score of 8 and the bottom drawing a score of 11. The far-right blocks contain drawings from participants in the Control Condition. The top drawing received a score of 7 and the bottom drawing a score of 5. Scores reflect the number of objects identified in drawings that were in the testing room photographs, which were used as references for raters (see Materials and Methods).

#### Stress Reactivity Assays

##### Cortisol

Salivary cortisol was evaluated at five timepoints: Baseline (upon arriving at the lab), Pre-Task (prior to task instructions), Immediate Post-Task (following TSST or control task), Delayed Post-Task (~15 mins post-task), or Recovery (~1 hr post-task). To avoid contaminating cortisol samples, we asked participants to refrain from physical activity, eating, drinking liquids other than water, smoking, and brushing their teeth during the 2-hrs prior to the study session, as well as to refrain from drinking water for at least 15 mins prior to the study session. After each study session, collection vials were capped, frozen and sent to Salimetrics, LLC (State College, PA) for cortisol assays. Following Salimetrics’ standard analysis procedures, all samples were measured twice and the average of the two values was used.

##### Psychophysiological data acquisition

Psychophysiological data were continuously recorded during the TSST and control task using two electrodes attached to the participants non-dominant palm (galvanic skin conductance level) and pulse oximetry to the non-dominate index finger (heart rate). Psychophysiological data were continuously recorded during the stress manipulation using a BIOPAC MP150 data acquisition system and EDA100C and OXY100E hardware (BIOPAC Systems Inc., Santa Barbara, CA) in conjunction with *AcqKnowledge* software at a sampling rate of 1000 Hz. Skin conductance level (SCL; measured in microSiemens [μS]) was recorded using two disposable dry Ag/AgCl laminated foam snap electrodes gelled with 0.5%-NaCl electrode gel and affixed to the thenar and hypothenar surfaces of the non-dominant palm. Heart rate in beats-per-minute was collected by affixing a finger clip pulse oximeter to the non-dominate index finger.

##### Psychophysiological data processing and analysis

Psychophysiological data were processed and analyzed using custom MATLAB scripts (Mathworks, Natick, MA). First, to minimize heart rate artifact, heart rate values exceeding 2 standard deviations beyond the mean or 200 beat-per-minute were considered artifact and interpolated to maintain a continuous signal. To remove high-frequency phasic responses and noise, the skin conductance level (SCL) and heart rate (HR) traces were low-pass filtered using a first-order bidirectional Butterworth Filter with a cut-off frequency of 5 Hz (Bach et al., 2013), smoothed using a 20 s moving window, and finally signal magnitudes were averaged into one-minute intervals (i.e., one SCL/HR measure per minute).

##### Stress summary variables

Cortisol reactivity (ng/dL; Δ Cortisol) was measured by subtracting baseline cortisol (Pre-Task sample) from the higher of the two post-stressor cortisol values (i.e. immediately or 15-mins post-stressor; Kim et al., 2019). We opted to use this metric as compared to other standard metrics (e.g. Area Under the Curve) due to imbalances in sample size of usable cortisol data across the collection timepoints. Psychophysiological stress responsivity was measured as the difference in heart rate levels (beats per minute; Δ HR) and SCL (μS; Δ SCL) during the first minute (typically just before the speech began) compared to the peak value across the binned time course (see above). Cortisol, SCL and HR have been identified as robust markers of stress responsivity to the TSST (see Allen et al., 2014 for review).

#### Sleep Monitoring

That evening, participants slept overnight in the sleep laboratory at Boston College, while being monitored with polysomnography. Polysomnographic recordings included electrooculography, chin electromyography, and electroencephalography (EEG; sampling rate = 200 Hz) from six scalp electrodes (F3, F4, C3, C4, O1, O2), each referenced to contralateral mastoid electrodes (M1, M2).

#### Free Recall Test

Twenty-four hours after stress manipulation, memory for objects in the testing room was assessed by free recall. Participants created drawings of the testing room, prompted by experimenters that they should include “as much detail possible”. Recent work demonstrates that drawings provide multidimensional episodic memory information (e.g. detailed spatial and object information) and provide insight into the form in which content is recalled (Bainbridge et al., 2019). Drawings were scored by 3 research assistants unassociated with data collection, who were blinded to the participants’ condition designation. Raters identified and counted the number of identifiable objects from drawings that were actually in the testing room using reference photographs (see **Figure 1**). This scoring was conducted once by each rater for each participant. Scoring was completed by consensus; if two raters disagreed on an object, the third rater broke the tie. All three judges agreed for 61% of identified objects and at least two judges agreed for 82% of identified objects. Free recall memory performance was computed as the total number of correctly identified testing room objects. For those in the Stress Condition only, we calculated separate scores for central objects (i.e. objects on the judges’ table or that were essential for evaluation) and peripheral objects (i.e. objects peripheral to the judges table that were non-evaluative). The central/peripheral distinction was not relevant for the Control Condition (i.e. evaluative objects were not present in this condition) and thus was not calculated.

#### Sleep EEG Analysis

##### Sleep Staging

NREM (N1, N2, and N3) and REM sleep was staged in 30-s epochs using American Academy of Sleep Medicine criteria (Iber et al., 2007) by a trained research specialist.

##### Artifact Rejection

EEG artifacts were automatically rejected using the Luna toolbox (Purcell et al., 2017, http://zzz.bwh.harvard.edu/luna/). For every epoch, the root mean square (RMS) and three Hjorth parameters (activity, mobility, and complexity) were calculated separately for each of the six EEG electrodes. Any epoch where at least one electrode exhibited an RMS or Hjorth parameter value > 3 SD from that electrode’s mean value was considered an artifact (Purcell et al., 2017). Subsequent analyses were performed on the artifact free data averaged across frontal (F3, F4) and central (C3, C4) electrodes.

##### Spectral Analysis

Power spectral density (PSD) was calculated using the pwelch function in MATLAB (Hamming window, 50% overlap). Power in the slow oscillation (0.3-1Hz), delta (1-4 Hz) and sigma (11-15Hz) bands were assessed in NREM sleep and theta power (4-7Hz) in REM sleep. Sleep spindles (~13.5 Hz) were detected in NREM sleep. To counteract the typical 1/f scaling, PSD was calculated not on the data time series itself, but on the temporal derivative of the data (Cox et al., 2017).

##### Sleep Spindle Detection

Sleep spindles were detected in N2 and N3 using a wavelet-based automated detector (Wamsley et al., 2012). The raw EEG signal underwent a time-frequency transformation using complex Morlet wavelets. Spindles were detected by applying a thresholding algorithm to the extracted wavelet scale with a center frequency of 13.5Hz. A spindle was detected whenever the wavelet signal exceeded a threshold of nine times the median signal amplitude of artifact-free epochs for at least 400ms (Mylonas et al., 2019).

#### Stress, Sleep and Memory Analysis

Group effects were evaluated with parametric statistics (i.e. analysis of variance, t-tests) or non-parametric tests, where indicated. Correlations were conducted using Pearson’s correlation coefficients or Spearman’s Rho, where appropriate. Fisher r-to-z transformation assessed group differences in correlation coefficients. Within each group, stress reactivity measures, including ΔCortisol, ΔHR, ΔSCL, and sleep stage percentages, including N2, N3, NREM (combined N2 and N3), and REM were independently correlated with memory performance. We then ran exploratory correlations between memory performance and NREM slow oscillation and delta power, sleep spindle number and density (spindles/min of NREM sleep) and REM sleep theta power (4-7 Hz). Values for each variable that exceeded 3 standard deviations above or below the within-group mean were excluded from analyses. Normality for each variable was tested using Shapiro-Wilk tests. Analyses were conducted in SPSS 25 unless otherwise indicated.

##### Interactive Effects of Stress and Sleep on Memory

We conducted a preliminary, exploratory analysis on the interactive effects of stress and sleep metrics on memory using multiple linear regression. Memory outcomes included total object memory and, in stressed participants only, central and peripheral object memory. Stress variables were ΔCortisol, ΔHR and ΔSCL. Sleep variables were NREM% and REM% sleep. Each tested model contained one stress variable and one sleep variable. Based on prior findings from Kim et al. (2019), we additionally ran a model in stressed participants that aimed to predict central object memory from ΔCortisol and REM sleep theta (θ) power.

#### Post-hoc power analysis

We conducted post-hoc power analyses using G*Power 3.1 based on our collected sample size and α = .05 for three effect sizes (small, medium and large). This was important to determine the limitations of our findings as this was a secondary analysis of a larger study. For group comparisons using t-tests, we had 12%, 51% and 89% power to detect small (d=.20), medium (d=.50) and large (d=.80) effects, respectively. For the non-parametric, within Stress Condition analysis of central compared to peripheral objects, we had 13%, 49% and 85% power to detect small (r=.10), medium (r=.30) and large (r=.50) effects, respectively. For analysis of variance (ANOVA), we had 5%, 20% and 82% power to detect small (partial η^2^ = .01), medium (partial η^2^ =.06) and large (partial η^2^ = .14) effects, respectively. For correlations, we had 7%, 21% and 48% power to detect small (r = .10), medium (r = .30) and large (r = .50) effects, respectively. Lastly, for multiple linear regressions we had 12%, 56% and 90% power to detect small (f^2^ = .02), medium (f^2^ = .15) and large (f^2^ = .35) effects, respectively in the Stress Condition. In the Control Condition, we had 12%, 58% and 91% power to detect small (f^2^ = .02), medium (f^2^ = .15) and large (f^2^ = .35) effects, respectively.

## Results

### Psychometrics and Sleep

Stressed and Control participants showed no significant differences in depression (BDI-II) or anxiety (BAI), objective (measured by actigraphy) and subjective (PSQI, MEQ, ESS) baseline sleep and post-encoding PSG-recorded sleep architecture (see **Table 1**).

### Stress reactivity

To evaluate task-associated cortisol curves, a 2 Condition (Stress, Control) by 5 Timepoint (Baseline, Pre-task, Immediate Post-Task, Delayed Post-Task, or Recovery) repeated measures ANOVA (Greenhouse-Geisser corrected) revealed significant main effects of Condition, F(1,43)=5.84, p=.02, partial η^2^ = .12, and Timepoint, F(1.75, 75.30)=4.28, p=.02, partial η^2^ = .09, and a significant Condition by Timepoint interaction, F(1.75, 75.30)=6.73, p=.003, partial η^2^ = .14 (see **Figure 2A**). Post-hoc t-tests revealed that participants in the Stress Condition showed markedly higher cortisol levels during the Delayed Post-Test Sample (~15-30 mins post-stressor, conforming to the typical peak in post-stressor cortisol) and marginally higher cortisol levels in the Immediate Post-Test sample. Cortisol levels did not significantly differ at other timepoints (see **Table 2**). Stressed participants exhibited significantly greater baseline to peak rises in stress reactivity for ΔCortisol, t(34.97) = 3.81, p = .001, d = 1.00, ΔSCL, t(56) = 2.05, p = .045, d= .054, and Δ HR, t(35.87) = 5.05, p<.001, d = 1.30 (see **Figure 2B-D**).

**Table 2.**
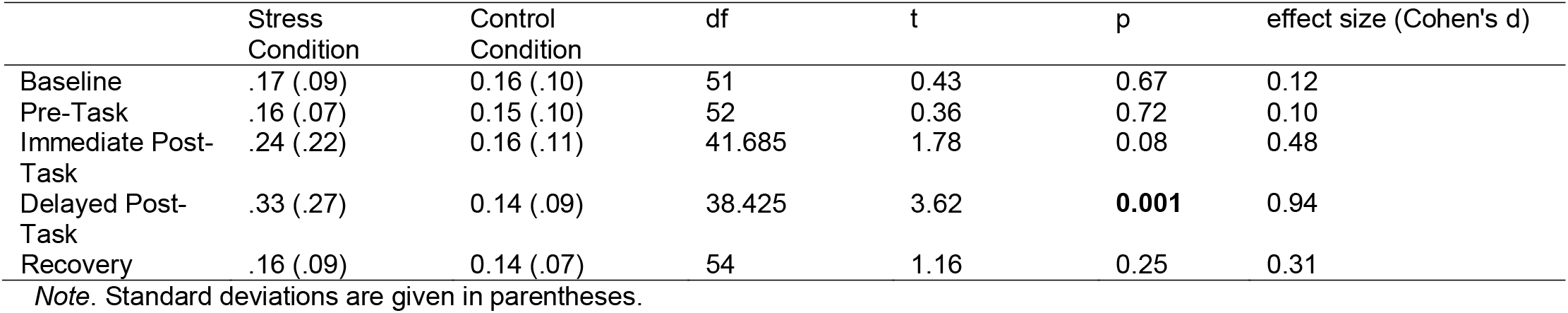
Mean cortisol levels across the experimental testing session.

|  | Stress<br>Condition | Control<br>Condition | df | t | p | effect size (Cohen's d) |
| --- | --- | --- | --- | --- | --- | --- |
| Baseline | .17 (.09) | 0.16 (.10) | 51 | 0.43 | 0.67 | 0.12 |
| Pre-Task | .16 (.07) | 0.15 (.10) | 52 | 0.36 | 0.72 | 0.10 |
| Immediate Post-Task | .24 (.22) | 0.16 (.11) | 41.685 | 1.78 | 0.08 | 0.48 |
| Delayed Post-Task | .33 (.27) | 0.14 (.09) | 38.425 | 3.62 | **0.001** | 0.94 |
| Recovery | .16 (.09) | 0.14 (.07) | 54 | 1.16 | 0.25 | 0.31 |

*Note.* Standard deviations are given in parentheses.

**Figure 2.**
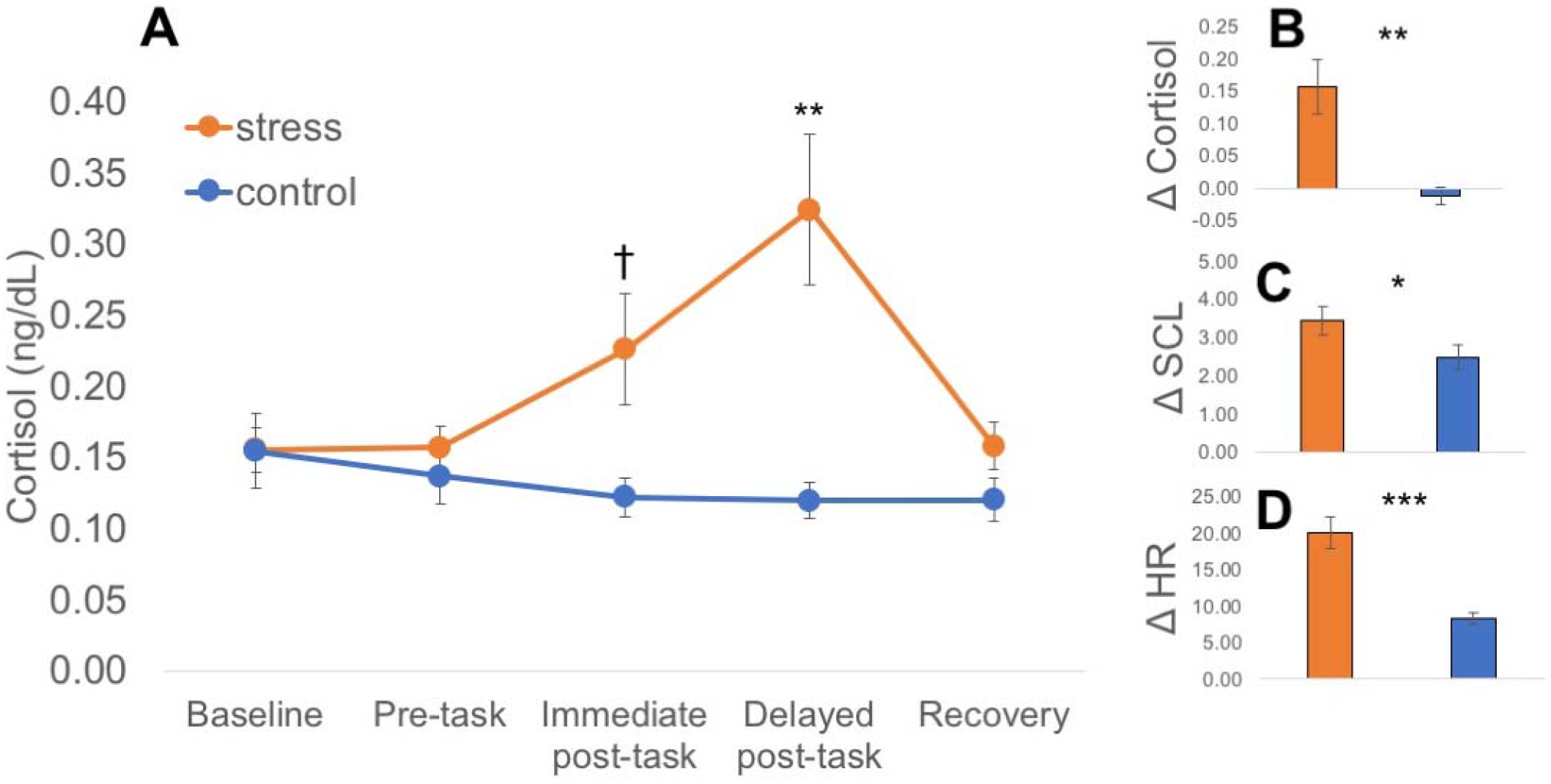
Stress reactivity plots. In (A) cortisol reactivity is plotted by study phase for Stress and Control participants. Plots (B-D) show the change in the metric from baseline to peak reactivity in Stress and Control participants. † p<.10, *p<.05, **p=.001, ***p<.001.

### Memory Performance and Stress

Compared to controls, stressed participants recalled significantly more objects from the testing environment (M_stress_=7.3. ± 1.9, M_control_=5.0 ± 1.8, t(56)=4.61, p<0.001, d=1.21) (see **Figure 1** for representative drawings and reference images), which was unsurprising because there were more objects present in the room in the stress relative to the Control Condition (e.g. camcorder, microphone, judge’s clipboards). When those evaluative objects were removed from analysis, group differences no longer reached significance (M_stress_ = 4.7 ± 1.5, M_control_ = 5.0 ± 1.8, t(56) = −.604, p = .548, d = .16). For those in the Stress Condition, the number of central objects remembered was greater than the number of peripheral objects remembered (Wilcoxon Signed-Rank Test: Mdn_central_ = 5, Mdn_peripheral_ = 2, Z = −4.57, p < 0.001, r = −3.0, 95% CI [−3.5, −2.0], d = 1.68), though there were many more peripheral objects that could have potentially been encoded and later remembered. This conceptually replicates prior work using a similar memory paradigm (Herten, Otto, et al., 2017; Herten, Pomrehn, et al., 2017; Wiemers et al., 2013, 2014). However, we will make no further interpretations of this finding given that the central/peripheral distinction for object memory could not be made for Control participants. Contrary to our hypothesis, ΔCortisol (r= −.093, p=.624), ΔSCL (r=.012, p=.949) and ΔHR (r=.20, p=.289) did not significantly correlate with central object memory performance in the Stress Condition. Likewise, the association between ΔSCL and total object memory (r = −.253, p = .185), likely driven by peripheral object memory (r = −.253, p = .185), was not significant and no other associations between memory performance (i.e. total object memory or peripheral object memory) and stress metrics were observed in this group (all ps>.419). In the Control Condition, ΔCortisol (r = −.207, p = .410), Δ SCL (r = −.158, p = .450) and ΔHR (r = −.049, p = .813) did not significantly correlate with total object memory.

### Correlations Between Memory Performance and Sleep

No significant associations were observed between memory and sleep metrics in the Stress Condition (*ps* > .219). In the Control Condition, total object memory was positively correlated with NREM sleep percentage (NREM%), r = .389, p = .041, NREM spindle number, r = .489, p = .01, NREM spindle density, r = .427, p = .026, and negatively correlated with REM sleep percentage (REM%), r = −.535, p = .003 (likely due to the inverse relationship between NREM and REM when quantified as a percentage of total sleep time; see **Figure 3)**. Further, these correlations consistently differed between groups, being significantly stronger in the Control Condition than the Stress Condition (Z_NREM%_ = 1.82, p= .034; Z_spindle number_= 1.52, p= .064; Z_spindle density_= 1.52, p= .065; Z_REM%_ = −2.57, p=.005). Memory was not correlated with any additional measure of sleep in controls (ps>.15).

**Figure 3.**
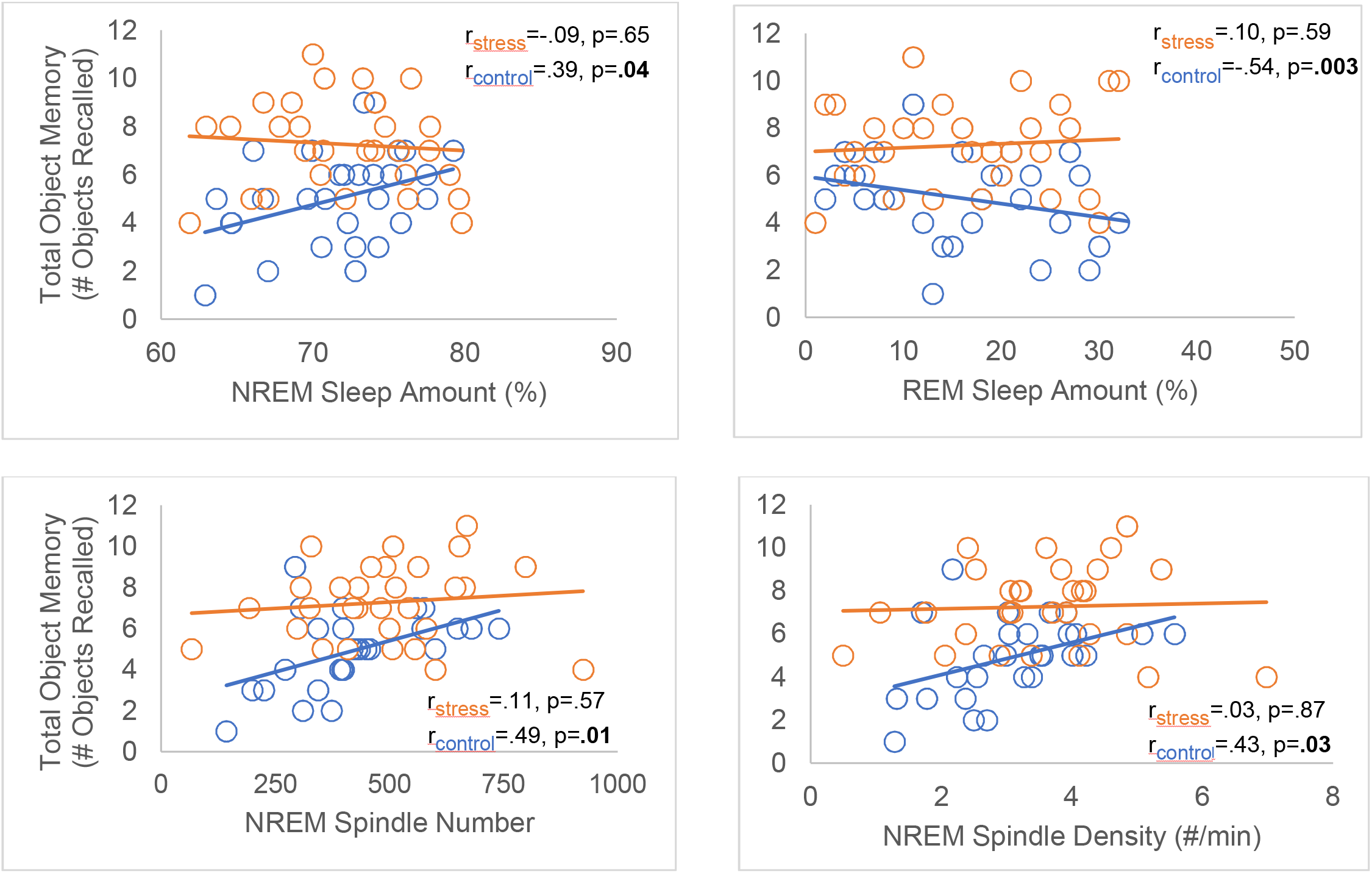
Correlations between memory performance and sleep physiology. Values and results from Stress participants are colored orange while those for Controls are colored blue.

### Interactions Between Stress, Sleep and Memory Performance

We did not observe any significant interactions between stress and sleep physiology on object memory, though we did observe a number of statistical trends (see **Tables 3-8** for regression model outputs). In Control participants, we observed an ΔSCL* NREM% interaction that predicted total object memory, β = −.38, 95% CI = [−20.26, .07], p = .051. At relatively low levels of NREM sleep, there was a positive association between skin conductance rise and memory performance, while at relatively higher levels of NREM sleep the opposite was true. For stressed participants, we observed a similar ΔCortisol* REM% interaction that predicted total object memory, β = −.47, β, 95% CI = [−98.77, 1.30], p = .056. At relatively low levels of REM sleep, cortisol rise was slightly negatively associated with memory performance while at relatively higher amounts of REM sleep there was an absence of association between cortisol rise and memory performance.

**Table 3.** Regression models for NREM sleep percentage in Control Condition.

| DV: Total Object Memory | Model 1 | $F(3, 17) = 2.12, p = .144, R^2 = .16$ | | |
| --- | --- | --- | --- | --- |
| | | | $\beta$ [95% CI] | $p$ |
| | | ΔCortisol | -.16 [-19.28, 10.45] | .513 |
|  |  | NREM% | .46 [-1.89, 47.54] | .068 |
| | | ΔCortisol* NREM% | -.14 [-388.21, 229.55] | .590 |
| | Model 2 | $F(3, 24) = 3.96, p = .022, R^2 = .270$ | | |
| | | | $\beta$ [95% CI] | $p$ |
| | | ΔSCL | -.26 [-.69, .13] | .176 |
|  |  | NREM% | .41 [1.84, 31.24] | .029 |
|  |  | **ΔSCL\* NREM%** | **-.38 [-20.26, .07]** | **.051** |
| | Model 3 | $F(3, 25) = 2.26, p = .110, R^2 = .131$ | | |
| | | | $\beta$ [95% CI] | $p$ |
| | | ΔHR | -.12 [-.23, .12] | .528 |
|  |  | NREM% | .45 [2.13, 35.90] | .029 |
| | | ΔHR* NREM% | -.13 [-5.55, 2.80] | .502 |

Note. CI = Confidence intervals, $R^2$ = Adjusted $R^2$ , NREM = Non-Rapid Eye Movement Sleep (NREM 2 + 3), SCL = Skin Conductance Level, HR = Heart Rate. Trending results bolded for emphasis.

**Table 4.** Regression models for REM sleep percentage in Control Condition.

| DV: Total Object Memory | Model 1 | $F(3, 17) = 2.09, p = .148, R^2 = .16$ | | |
| --- | --- | --- | --- | --- |
| | | | $\beta$ [95% CI] | $p$ |
| | | ΔCortisol | -.21 [-20.08, 7.94] | .368 |
|  |  | REM% | -.53 [-61.41, 5.41] | .094 |
| | | ΔCortisol* REM% | -.03 [-349.80, 321.62] | .930 |
| | Model 2 | $F(3, 24) = 5.01, p = .009, R^2 = .33$ | | |
| | | | $\beta$ [95% CI] | $p$ |
| | | ΔSCL | -.22 [-.62, .140] | .204 |
|  |  | REM% | -.53 [-36.13, -7.04] | .006 |
| | | ΔSCL* REM% | .25 [-2.73, 16.69] | .150 |
| | Model 3 | $F(3, 25) = 3.60, p = .030, R^2 = .24$ | | |
| | | | $\beta$ [95% CI] | $p$ |
| | | ΔHR | .07 [-.14, .20] | .711 |
|  |  | REM% | -.58 [-40.30, -8.33] | .005 |
| | | ΔHR* REM% | -.002 [-4.00, 3.97] | .993 |

Note. CI = Confidence intervals, $R^2$ = Adjusted $R^2$ , REM = Rapid Eye Movement, SCL = Skin Conductance Level, HR = Heart Rate.

**Table 5.**
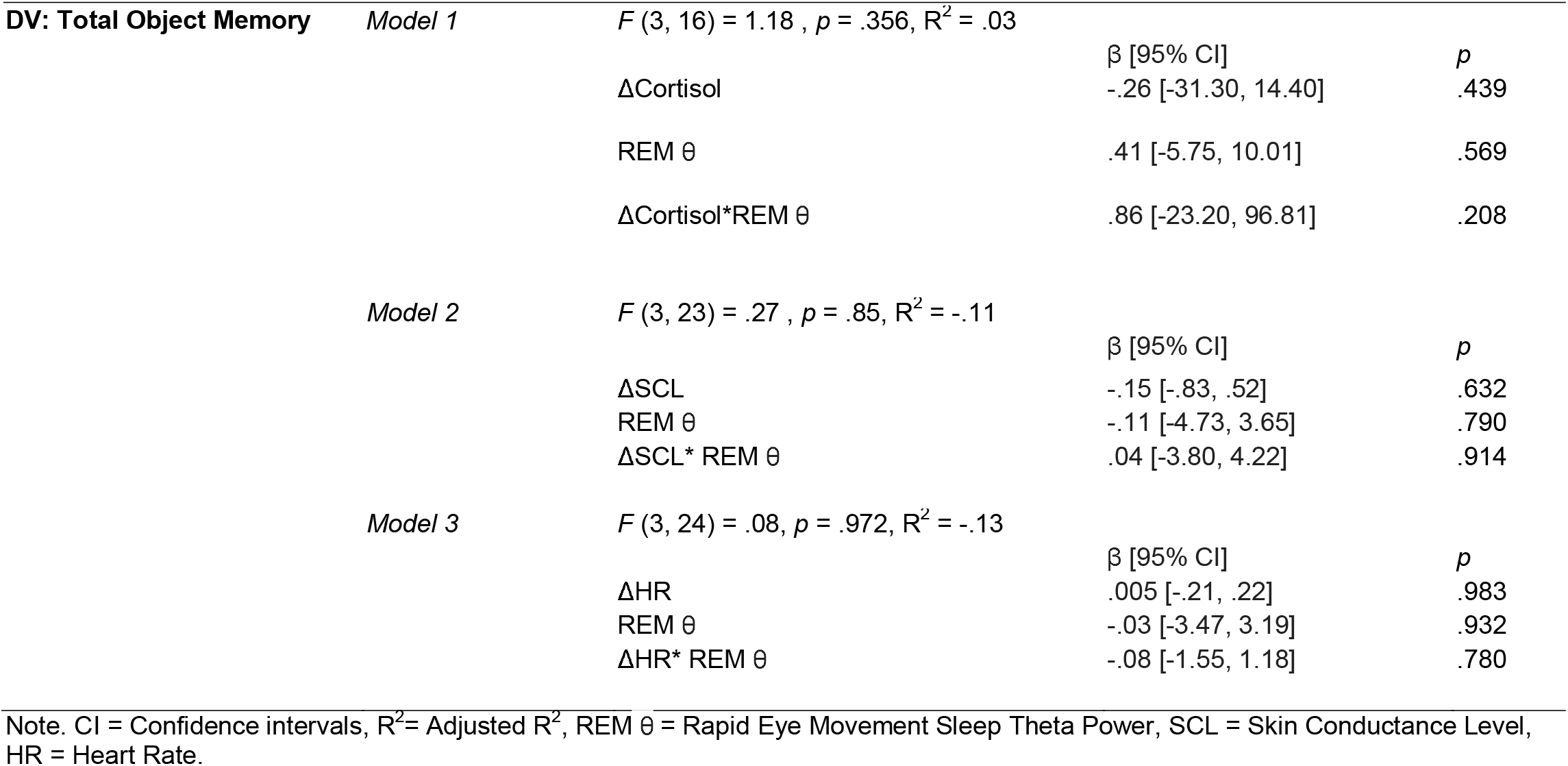
Regression models for REM sleep theta power (REM θ) in Control Condition.

|  |  |  |  |  |
| --- | --- | --- | --- | --- |
| **DV: Total Object Memory** | *Model 1* | $F(3, 16) = 1.18, p = .356, R^2 = .03$ | | |
| | | | $\beta$ [95% CI] | $p$ |
| | | ΔCortisol | -.26 [-31.30, 14.40] | .439 |
| | | REM $\theta$ | .41 [-5.75, 10.01] | .569 |
| | | ΔCortisol*REM $\theta$ | .86 [-23.20, 96.81] | .208 |
| | *Model 2* | $F(3, 23) = .27, p = .85, R^2 = -.11$ | | |
| | | | $\beta$ [95% CI] | $p$ |
| | | ΔSCL | -.15 [-.83, .52] | .632 |
| | | REM $\theta$ | -.11 [-4.73, 3.65] | .790 |
| | | ΔSCL*REM $\theta$ | .04 [-3.80, 4.22] | .914 |
| | *Model 3* | $F(3, 24) = .08, p = .972, R^2 = -.13$ | | |
| | | | $\beta$ [95% CI] | $p$ |
| | | ΔHR | .005 [-.21, .22] | .983 |
| | | REM $\theta$ | -.03 [-3.47, 3.19] | .932 |
| | | ΔHR*REM $\theta$ | -.08 [-1.55, 1.18] | .780 |

Note. CI = Confidence intervals, $R^2$ = Adjusted $R^2$ , REM $\theta$ = Rapid Eye Movement Sleep Theta Power, SCL = Skin Conductance Level, HR = Heart Rate.

**Table 6.**
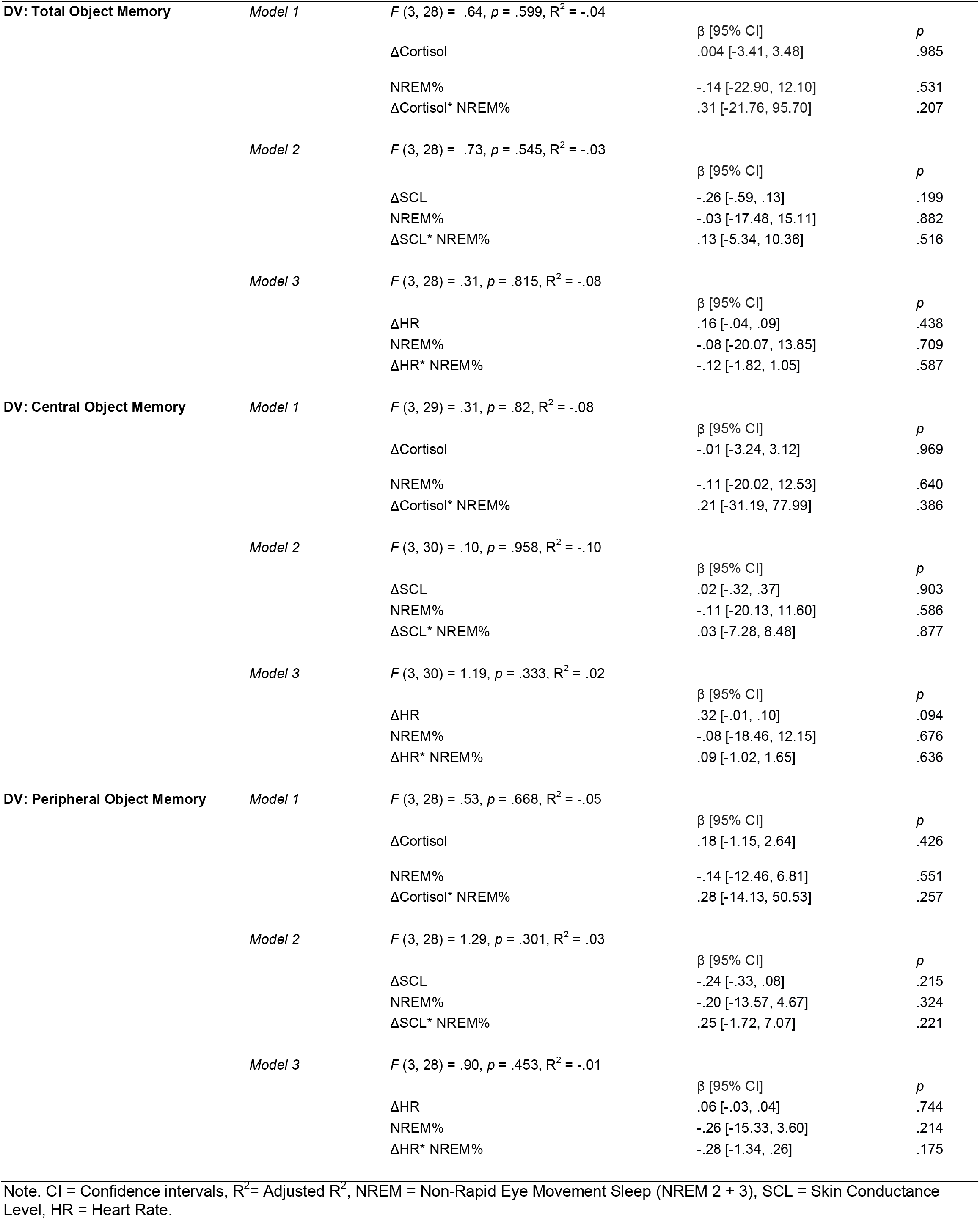
Regression models for NREM sleep percentage in Stress Condition

**Table 7.**
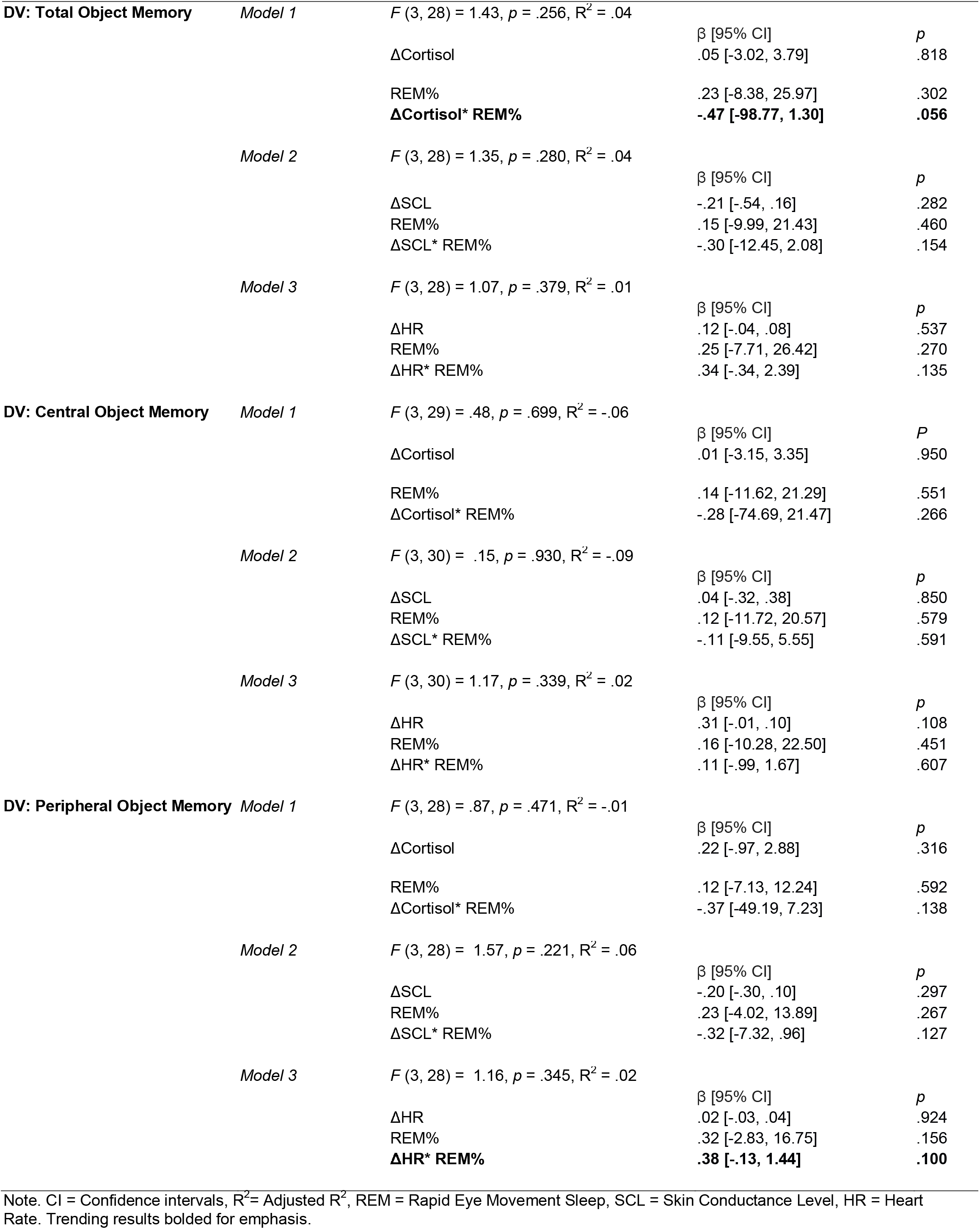
Regression models for REM sleep percentage in Stress Condition

**Table 8.**
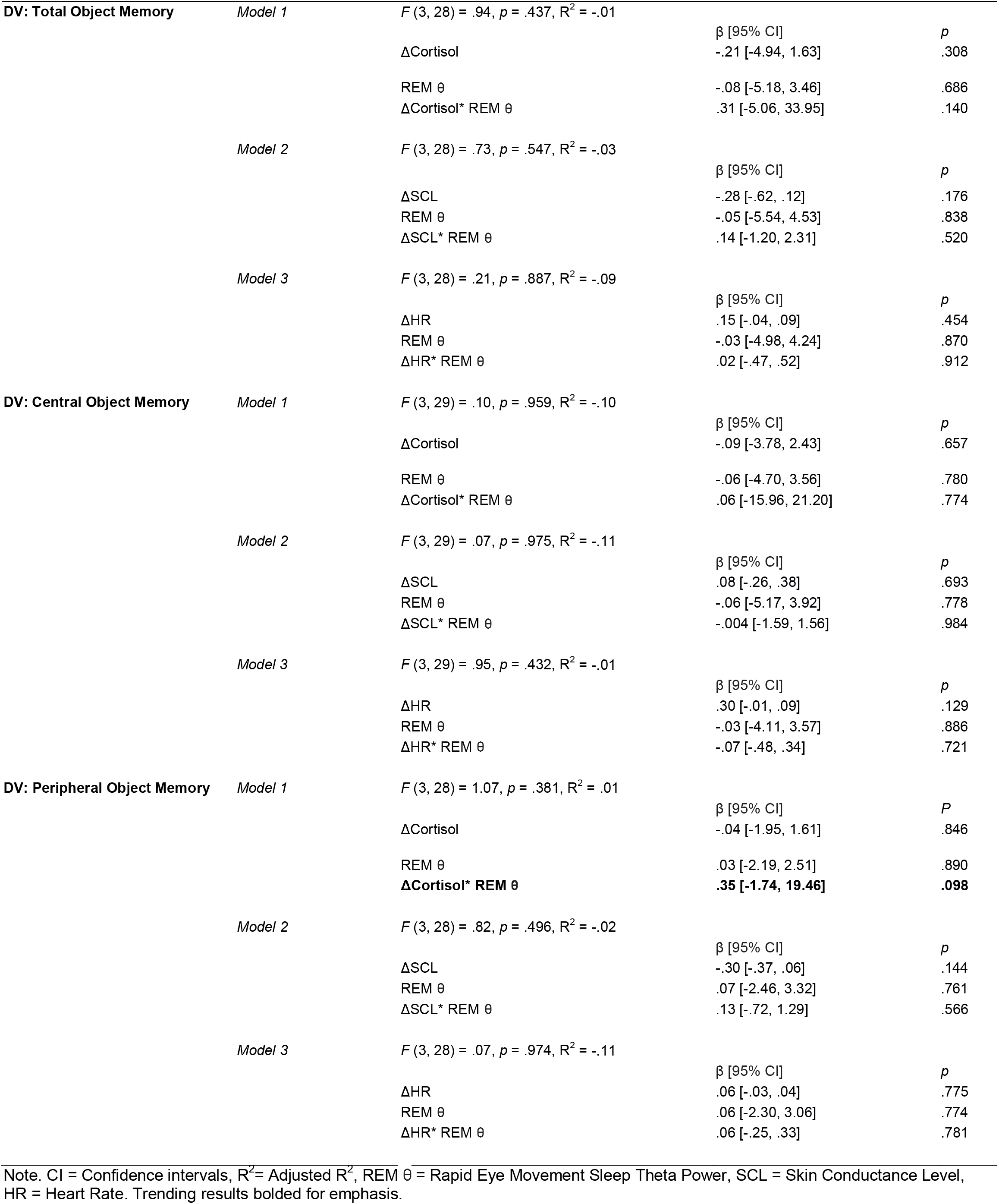
Regression models for REM sleep theta power (REM θ) in Stress Condition.

## Discussion

Contrary to our hypothesis, but in alignment with prior reports using a similar paradigm (Wiemers et al., 2013), we did not observe strong evidence for a relationship between stress response magnitude during a psychosocial stressor and memory performance. It may be that neuromodulators associated with stress responsivity may only need to reach a critical threshold to “tag” salient information for selective long-term storage (Payne & Kensinger, 2018), though future work is necessary to substantiate this hypothesis.

Also contrary to our prediction, stress responsivity and sleep architecture did not significantly interact to influence memory performance. Though it is notable that our sample sizes are comparable or exceed those of other similar studies in this research area, we interpret such null findings with caution as some may have resulted from a lack of statistical power. In fact, when these same participants had their memory examined for emotional pictures presented soon after the stressor, Kim et al. (2019) found an association between REM sleep and positive picture recall in participants who mounted high cortisol responses to the TSST. This combination of results suggests the intriguing possibility that information encountered *soon after* a stressor may be more affected by sleep than memory encountered *during* a stressor. Although future work will be needed to address this possibility directly.

Our findings that NREM sleep physiology correlated with object memory in non-stressed participants aligns with a large body of literature suggesting NREM sleep physiology impacts episodic memory in controlled laboratory tasks (For review see Rasch & Born, 2013). Our findings suggest that NREM’s faciliatory effect on episodic memory consolidation may hold in real-world situations.

While stress exposure did not impact sleep macroarchitecture (i.e. sleep stage percentages or total sleep time), it is possible that subtle autonomic measures during sleep could better predict subsequent memory. For example, Groch et al. (2011) demonstrated that norepinephrine blockade with clonidine during sleep did not affect sleep macroarchitecture, but did diminish emotional memory biases compared to a placebo condition. Conversely, stress can fragment sleep leading to increases in stress neuromodulators (Âkerstedt, 2006) which might then paradoxically enhance memory, albeit selectively for emotional traces. While we did not measure autonomic activity during post-stressor sleep, implementation of such metrics in future work may clarify subtle changes to sleep that impact long-term memory retention.

### Limitations

Several limitations exist in the current study. First, based on post-hoc power analyses, our study could only reliably detect large effects. Follow-up work will require larger sample sizes to verify our current findings. Due to constraints imposed by the task design of the overarching study from which this is a secondary analysis, there were imbalances in the number of objects available for Stress and Controls participants to encode. Specifically, the testing environment included two salient evaluation, a camcorder and microphone, and several subtler evaluation objects including clipboards that judges recording their assessments on and a stopwatch used to time the speech and arithmetic task. Without these, a comparable central/peripheral memory trade-off could not be established for Controls. The original intention for excluding these objects from the room for Control participants was to minimize confounding evaluative stress their presence may have induced. These objects may also have disproportionately captured the attention of the Stress Condition participants, shifting attention away from peripheral objects and thus resulting in poor associations between physiological metrics and memory. However, at least one study, which used a similar task paradigm, concluded that object fixation time during encoding did not mediate the relationship between stress and memory performance (Herten, Otto, et al., 2017). Future work may benefit from including similar objects for Controls, though manipulate instructions to participants to de-valance these objects (e.g. “These objects are present for all participants, but are not recoding you). This study did not directly compare a sleep delay to wake delay condition, which has been standard in prior studies investigating stress and sleep’s impact on memory (e.g. (Bennion et al., 2015; Cunningham et al., 2014). We therefore cannot completely rule out the possibility that stress and sleep interact in a way that we could not quantify with individual differences measures alone.

## Conclusions

Our findings do not support an interactive role of stress and sleep physiology on naturalistic episodic memories encoded during a stressful experience. However, in the absence of social stress induction, next-day memory for testing room details was associated with NREM sleep obtained the night after encoding. The latter finding expands upon the growing literature implicating NREM sleep physiology in memory consolidation by using a task the better approximates a real-life encoding experience. Additional controls and expanded samples size may be necessary to confirm our current pattern of findings.

## Acknowledgments

The authors thank Stephanie Sherman, Kevin Fredericks, Lauren Liu, Tala Berro, Haley DiBiase, Ross Mair, Tammy Moran, Olivia Hampton, Sabie Marcellus, Christopher Bischoff, Spurti Vemuri, Nicholas Pontillo, Haley Woloshen, Thor Janson, and Yasmin Yacoby for their help with data collection; Claire Cushman, Isabella Turco and Michael Frank for their help with coding participant drawings; and thanks to SomnoSure, Inc. for scoring the sleep PSG records.

## Data Availability

Data are accessible on Open Science Framework:

- Bottary, R. (2021, January 27). Investigation of the Stress and Sleep Physiology Correlates of Next-Day Memory for Details of a Social Stressor Testing Environment. Retrieved from osf.io/gny3k.

## References

Âkerstedt, T. (2006). Psychosocial stress and impaired sleep. Scandinavian Journal of Work, Environment & Health, 32(6), 493–501. JSTOR.

Allen, A. P., Kennedy, P. J., Cryan, J. F., Dinan, T. G., & Clarke, G. (2014). Biological and psychological markers of stress in humans: Focus on the Trier Social Stress Test. Neuroscience & Biobehavioral Reviews, 38, 94–124. https://doi.org/10.1016/j.neubiorev.2013.11.005

Bach, D. R., Friston, K. J., & Dolan, R. J. (2013). An improved algorithm for model-based analysis of evoked skin conductance responses. Biological Psychology, 94(3), 490–497. https://doi.org/10.1016/j.biopsycho.2013.09.010

Bainbridge, W. A., Hall, E. H., & Baker, C. I. (2019). Drawings of real-world scenes during free recall reveal detailed object and spatial information in memory. Nature Communications, 10(1), 5. https://doi.org/10.1038/s41467-018-07830-6

Beck, A., Steer, R., & Brown, G. (1996). Beck Depression Inventory-II. The Psychological Corporation, San Antonio, TX.

Bennion, K. A., Mickley Steinmetz, K. R., Kensinger, E. A., & Payne, J. D. (2015). Sleep and Cortisol Interact to Support Memory Consolidation. Cerebral Cortex, 25(3), 646–657. https://doi.org/10.1093/cercor/bht255

Birkett, M. A. (2011). The Trier Social Stress Test Protocol for Inducing Psychological Stress. Journal of Visualized Experiments◻: JoVE, 56. https://doi.org/10.3791/3238

Buysse, D. J., Reynolds, C. F., Monk, T. H., Berman, S. R., & Kupfer, D. J. (1989). The Pittsburgh sleep quality index: A new instrument for psychiatric practice and research. Psychiatry Research, 28(2), 193–213. https://doi.org/10.1016/0165-1781(89)90047-4

Cox, R., Schapiro, A. C., Manoach, D. S., & Stickgold, R. (2017). Individual Differences in Frequency and Topography of Slow and Fast Sleep Spindles. Frontiers in Human Neuroscience, 11. https://doi.org/10.3389/fnhum.2017.00433

Cunningham, T. J., Crowell, C. R., Alger, S. E., Kensinger, E. A., Villano, M. A., Mattingly, S. M., & Payne, J. D. (2014). Psychophysiological arousal at encoding leads to reduced reactivity but enhanced emotional memory following sleep. Neurobiology of Learning and Memory, 114, 155–164. https://doi.org/10.1016/j.nlm.2014.06.002

Groch, S., Wilhelm, I., Diekelmann, S., Sayk, F., Gais, S., & Born, J. (2011). Contribution of norepinephrine to emotional memory consolidation during sleep. Psychoneuroendocrinology, 36(9), 1342–1350. https://doi.org/10.1016/j.psyneuen.2011.03.006

Herten, N., Otto, T., & Wolf, O. T. (2017). The role of eye fixation in memory enhancement under stress – An eye tracking study. Neurobiology of Learning and Memory, 140, 134–144. https://doi.org/10.1016/j.nlm.2017.02.016

Herten, N., Pomrehn, D., & Wolf, O. T. (2017). Memory for objects and startle responsivity in the immediate aftermath of exposure to the Trier Social Stress Test. Behavioural Brain Research, 326, 272–280. https://doi.org/10.1016/j.bbr.2017.03.002

Horne, J. A., & Östberg, O. (1976). A self-assessment questionnaire to determine morningness-eveningness in human circadian rhythms. International Journal of Chronobiology, 4, 97–110.

Hutchison, I. C., & Rathore, S. (2015). The role of REM sleep theta activity in emotional memory. Frontiers in Psychology, 6. https://doi.org/10.3389/fpsyg.2015.01439

Iber, C., Ancoli-Israel, S., Chesson, A. L., & Quan, S. F. (2007). The AASM manual for the scoring of sleep and associated events: Rules, terminology and technical specifications (Vol. 1).

American Academy of Sleep Medicine Westchester, IL. Johns, M. W. (1991). A New Method for Measuring Daytime Sleepiness: The Epworth Sleepiness Scale. Sleep, 14(6), 540–545. https://doi.org/10.1093/sleep/14.6.540

Kim, S. Y., Kark, S. M., Daley, R. T., Alger, S. E., Rebouças, D., Kensinger, E. A., & Payne, J. D. (2019). Interactive effects of stress reactivity and rapid eye movement sleep theta activity on emotional memory formation. Hippocampus, 1–13. https://doi.org/10.1002/hipo.23138

Kirschbaum, C., Pirke, K.-M., & Hellhammer, D. H. (1993). The ‘Trier Social Stress Test’ – A Tool for Investigating Psychobiological Stress Responses in a Laboratory Setting. Neuropsychobiology, 28(1-2), 76–81. https://doi.org/10.1159/000119004

Latchoumane, C.-F. V., Ngo, H.-V. V., Born, J., & Shin, H.-S. (2017). Thalamic Spindles Promote Memory Formation during Sleep through Triple Phase-Locking of Cortical, Thalamic, and Hippocampal Rhythms. Neuron, 95(2), 424–435.e6. https://doi.org/10.1016/j.neuron.2017.06.025

Lehmann, M., Schreiner, T., Seifritz, E., & Rasch, B. (2016). Emotional arousal modulates oscillatory correlates of targeted memory reactivation during NREM, but not REM sleep. Scientific Reports, 6(1), 39229. https://doi.org/10.1038/srep39229

Loeffler, S. N., Hennig, J., & Peper, M. (2017). Psychophysiological Assessment of Social Stress in Natural and Laboratory Situations: Using the Experience Sampling Method and Additional Heart Rate Measures. Journal of Psychophysiology, 31(2), 67–77. https://doi.org/10.1027/0269-8803/a000170

Payne, J. D., & Kensinger, E. A. (2018). Stress, sleep, and the selective consolidation of emotional memories. Current Opinion in Behavioral Sciences, 19, 36–43. https://doi.org/10.1016/j.cobeha.2017.09.006

Purcell, S. M., Manoach, D. S., Demanuele, C., Cade, B. E., Mariani, S., Cox, R., Panagiotaropoulou, G., Saxena, R., Pan, J. Q., Smoller, J. W., Redline, S., & Stickgold, R. (2017). Characterizing sleep spindles in 11,630 individuals from the National Sleep Research Resource. Nature Communications, 8(1), 15930. https://doi.org/10.1038/ncomms15930

Rasch, B., & Born, J. (2013). About Sleep’s Role in Memory. Physiological Reviews, 93(2), 681–766. https://doi.org/10.1152/physrev.00032.2012

Shields, G. S., Sazma, M. A., McCullough, A. M., & Yonelinas, A. P. (2017). The effects of acute stress on episodic memory: A meta-analysis and integrative review. Psychological Bulletin, 143(6), 636–675. https://doi.org/10.1037/bul0000100

Steer, R. A., & Beck, A. T. (1997). Beck Anxiety Inventory. In Evaluating stress: A book of resources (pp. 23–40). Scarecrow Education.

Wamsley, E. J., Tucker, M. A., Shinn, A. K., Ono, K. E., McKinley, S. K., Ely, A. V., Goff, D. C., Stickgold, R., & Manoach, D. S. (2012). Reduced Sleep Spindles and Spindle Coherence in Schizophrenia: Mechanisms of Impaired Memory Consolidation? Biological Psychiatry, 71(2), 154–161. https://doi.org/10.1016/j.biopsych.2011.08.008

Wiemers, U. S., Sauvage, M. M., Schoofs, D., Hamacher-Dang, T. C., & Wolf, O. T. (2013). What we remember from a stressful episode. Psychoneuroendocrinology, 38(10), 2268–2277. https://doi.org/10.1016/j.psyneuen.2013.04.015

Wiemers, U. S., Sauvage, M. M., & Wolf, O. T. (2014). Odors as effective retrieval cues for stressful episodes. Neurobiology of Learning and Memory, 112, 230–236. https://doi.org/10.1016/j.nlm.2013.10.004

Wolf, O. T. (2019). Memories of and influenced by the Trier Social Stress Test. Psychoneuroendocrinology, 105, 98–104. https://doi.org/10.1016/j.psyneuen.2018.10.031

